# Interpreting citizen science data for larval surveillance: lessons from storm drain reports in Mosquito Alert

**DOI:** 10.64898/2026.11.23.768715

**Authors:** Ana Aguilar-Pineda, Isabella Jiménez-Jiménez, Fabio Sánchez

**Affiliations:** CIMPA–Escuela de Matemática, Universidad de Costa Rica, San José, Costa Rica; Colegio Científico de Costa Rica, Sede San Pedro, San José, Costa Rica

**Keywords:** Citizen science, larval surveillance, Mosquito Alert, storm drains, breeding sites, *Aedes albopictus*, *Aedes aegypti*, data visualization

## Abstract

Mosquito-borne diseases are an increasing public health concern, and urban drainage infrastructure such as storm drains and catch basins can provide breeding habitat for *Aedes* vectors, including *Aedes albopictus*. Detecting these sites over large areas is costly with traditional surveillance alone, and citizen science offers a complementary, lower-cost source of geolocated observations. Mosquito Alert is a citizen science platform, active since 2014, through which participants report adult mosquitoes, bites, and potential breeding sites. In this work, we present a descriptive analysis of the 17,880 storm drain breeding sites reported through the platform from June 2014 to August 2026, obtained from the project’s public map. We counted the reported sites by country and geographic coordinate, examined when reports are submitted and which fields are recorded, estimated how often water and larvae are reported, and mapped the records globally. We also developed an interactive map to help inspect regions where many records overlap. Spain accounts for 90% of the records, 82% of all reports were submitted between April and June, and half of them fall on only 50 days. Group events, defined as five or more users reporting in the same municipality on the same day, account for 63% of the records. Water was reported in 43% of the storm drains: 33% in group events and 61% in the remaining reports. Larvae information exists for only 1,739 records; almost all are drains with water, and 28.5% of those had larvae, with higher shares in the warmer months and where participants noticed adult mosquitoes nearby. Our aim is not to model mosquito populations but to describe the structure of the data and highlight patterns worth further investigation. Because citizen science reporting effort varies across space, time, and socioeconomic groups, we discuss how sampling bias and other limitations of volunteer-based data affect the interpretation of these patterns and what they imply for the use of such data in larval surveillance.

## 1 Introduction

Mosquito-borne diseases are a growing public health problem worldwide. Two of the most important vectors are *Aedes aegypti* and *Aedes albopictus*, commonly known as the yellow fever mosquito and the Asian tiger mosquito, respectively, which can transmit dengue, chikungunya, and Zika viruses. In endemic settings such as Costa Rica, dengue and chikungunya circulate through the same *Aedes* vectors [1], and dengue incidence closely follows climate and local environmental conditions [2, 3]. In recent decades, both species have expanded beyond their native ranges, mainly through international trade and travel, and are now distributed across multiple continents, including parts of Europe and North America [4]. In Europe, *Ae. albopictus* was introduced mainly through international trade in used tires and ornamental plants, and its subsequent spread has raised public health concerns [5].

The risk it poses has already materialized. The first outbreak of autochthonous chikungunya transmission in continental Europe occurred in north-eastern Italy in 2007, where *Ae. albopictus* was identified as the likely transmission vector [6]. Since then, several European countries have reported locally acquired cases of dengue and chikungunya [7]. In Spain, the species was first detected in 2004 in Sant Cugat del Vallès, Catalonia [8]. Surveillance documented its spread along the Mediterranean coast and into the Balearic Islands [9].

Because *Aedes* larvae develop in water, much of vector control focuses on locating larval breeding sites and removing or treating them. In cities, this is challenging because many small containers and water-holding structures can provide suitable habitats. Storm drains and catch basins are a relevant example. Water can accumulate in these structures and create conditions suitable for mosquito development. A study conducted in Shanghai found that *Ae. albopictus* was the predominant mosquito species in stormwater catch basins and manhole chambers, with catch basins representing an important source of mosquito production [10]. This highlights the importance of including drainage infrastructure in larval surveillance. For effective control, it is important to identify these potential breeding sites and determine whether they contain stagnant water or mosquito larvae. However, collecting this information across large areas using traditional field surveillance alone can be costly and labor-intensive [11].

Citizen science has emerged as a complementary approach to traditional surveillance. Through smartphone applications, volunteers can submit geolocated observations and photographs that experts can assess and validate. These platforms can provide relatively low-cost, large-scale, and timely information for mosquito surveillance [11, 12]. Mosquito Alert is a citizen science platform launched in 2014, through which participants can report adult mosquitoes, mosquito bites, and potential breeding sites [13]. In Spain, the platform has demonstrated its potential to provide early warnings of the spread of *Ae. albopictus*, complementing information obtained through traditional surveillance methods [12]. More recently, citizen science data from Mosquito Alert have contributed approximately one-third of the municipal detections of *Ae. albopictus* in Spain since the platform was launched [11]. Among the potential breeding sites reported by participants, storm drains are particularly relevant because they can support mosquito larval development, as demonstrated by studies of urban drainage systems [10]. The reports submitted through the platform are publicly accessible through the project’s online map [14].

However, citizen science data are not collected uniformly. Reporting effort can vary across geographical areas and socioeconomic groups, potentially introducing spatial sampling biases that affect the apparent distribution of observations [15]. Consequently, analyses based on these data should begin with a careful description of the spatial and temporal distribution of the available records, accounting for the limitations associated with volunteer-based reporting.

In this work, we present a descriptive analysis of storm drain breeding sites reported through Mosquito Alert. We obtained records from the project’s public map [14], selecting the two storm drain categories available in the application (storm drains with water and dry storm drains) from the start of the project in 2014 through August 2026. Our aim is not to model mosquito populations, but to describe the structure of the data and identify patterns that may be worth investigating further. To achieve this, we count reported breeding sites by country and geographic coordinate, describe when reports are submitted and which fields are recorded in each period, identify days on which many participants report in the same place, estimate how frequently water and larvae are reported, and map the records globally, paying particular attention to areas with the most reports. We also developed an interactive version of the map to facilitate the inspection of regions where multiple records overlap geographically.

The rest of the paper is organized as follows. First, we describe the data and the visualization methods used to produce the figures. Then, we present the results of each analysis and, finally, discuss their interpretation and limitations.

## 2 Methodology

### 2.1 Data Collection

Data were obtained from Mosquito Alert [12, 14], a citizen science platform designed for the research, surveillance, and management of disease-vector mosquitoes, specifically the Asian tiger mosquito (*Aedes albopictus*) and the yellow fever mosquito (*Aedes aegypti*). These species serve as vectors for arboviruses of global significance, including dengue, chikungunya, and Zika. The platform operates via a free mobile application that enables citizens to submit observational reports of adult mosquitoes, breeding sites, and bite frequency. Expert entomologists validate adult mosquito reports that include photographs by independently reviewing the images [13, 15]. We found no description of an equivalent validation for breeding-site reports, and the storm drain records analyzed here include no validation field. We therefore treat the information in these records, including the presence of water and larvae, as reported by the participants.

For this study, we retrieved all published storm drain reports from the Mosquito Alert public map on September 12, 2026. The dataset contains *N* = 17,880 records created between June 13, 2014, and August 16, 2026, submitted by 3,313 users in 66 countries. Most reports come from Spain, where the platform was launched, with smaller numbers from other countries such as Italy and Austria. The analysis used geographic coordinates (latitude and longitude), local creation time, country, municipality, an anonymous user identifier, and three yes/no fields: whether the storm drain contained water, whether larvae were observed, and whether the participant noticed adult mosquitoes nearby.

The yes/no fields are not equally complete (Table 1). The presence of water is recorded for every report. The presence of larvae is recorded for only 1,739 reports (9.7%), and its availability depends on the period and on the water field: it is present for most drains with water reported until 2020 and again from late May 2026, is essentially absent between 2021 and 2025, and is almost never recorded for dry drains. Any statement about larvae therefore refers to drains with water reported in those two periods. The presence of adult mosquitoes nearby is recorded for 2,502 reports, all but one of them submitted between 2015 and 2020.

**Table 1.** Number of storm drain reports by period and water status, and number of those reports with information on larvae.

|  |  | Drains with water |  | Dry drains |  |
|---|---|---|---|---|---|
| Period | Reports | Reports | Larvae recorded | Reports | Larvae recorded |
| 2014–2020 | 3,137 | 1,847 | 1,526 | 1,290 | 12 |
| 2021–2025 | 13,651 | 5,706 | 1 | 7,945 | 0 |
| 2026 | 1,092 | 203 | 199 | 889 | 1 |
| Total | 17,880 | 7,756 | 1,726 | 10,124 | 13 |

### 2.2 Analysis and Visualization

All analyses were carried out in Python. Records were filtered, grouped, and counted with pandas, and figures were produced with matplotlib. The continent outlines of the world maps come from Natural Earth (1:50m) and were loaded with Cartopy.

#### Counts by country and coordinate

Records were counted by country, and the 20 countries with the most records are shown in Figure 1a. To count records by location, latitude and longitude were rounded to three decimals, so that reports from almost the same place, within roughly 100 m, are assigned to the same coordinate; the 30 coordinates with the most records are shown in Figure 1b. To compare water and larvae, we excluded records with a missing value in either field, and counted the remaining records in each of the four combinations of water and larvae (Figure 1c). In these charts, bars are sorted from largest to smallest and labeled with their values.

**Figure 1.**
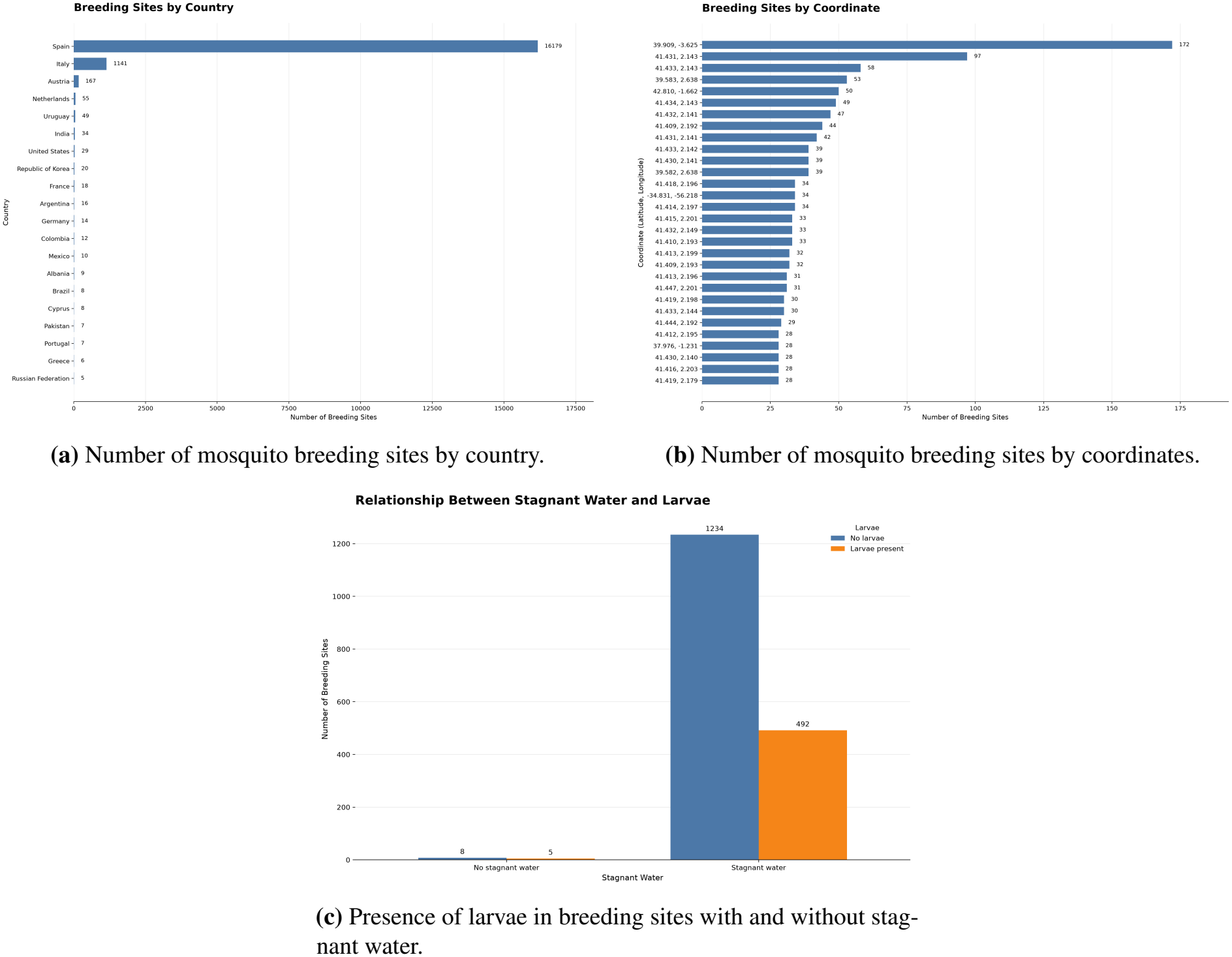
Summary of the storm drain breeding site records: (a) top 20 countries, (b) top 30 coordinates, and (c) stagnant water versus larvae.

#### World maps

For the world map of breeding sites (Figure 2), records were grouped by rounded coordinate. Each coordinate is drawn as a circle whose size and color increase with the number of records, from yellow to dark purple. Because a few coordinates have many more records than the rest, the color scale follows a power law, which keeps coordinates with few records visible. The map uses the Robinson projection, implemented in NumPy by interpolating its reference table, which is tabulated every 5^°^ of latitude. The ten countries with the most records are labeled with their totals, and three areas with many records are marked with boxes: the Iberian Peninsula (A), Barcelona (B), and Rome and its surroundings (C). The number of breeding sites and of unique coordinates in each area is given below the map. A second map of the same kind (Figure 7) includes only records with information on both water and larvae; there, circle size indicates the number of records at each coordinate, and color indicates the share with larvae.

**Figure 2.**
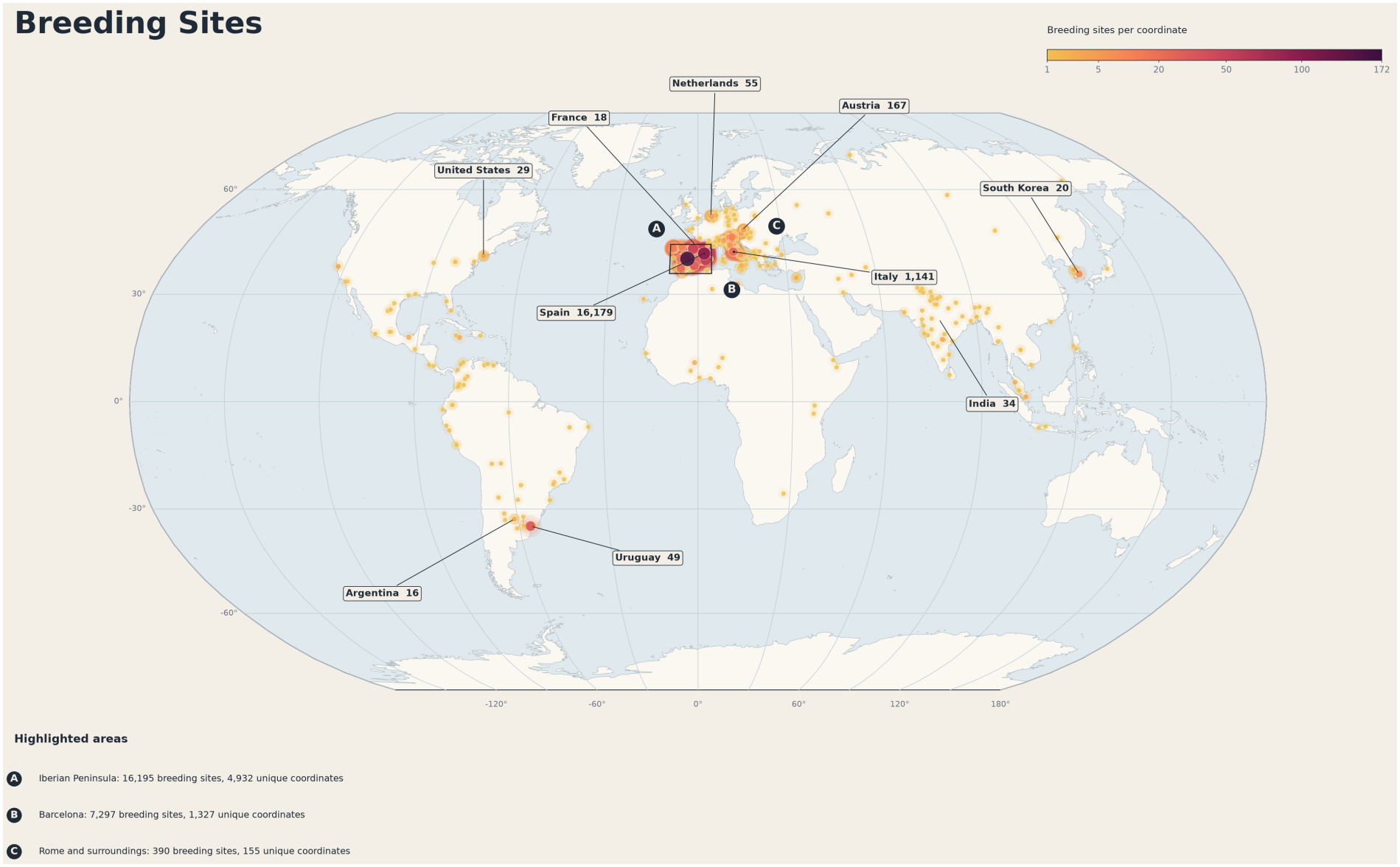
World map of breeding sites. Each circle represents a coordinate, with size and color indicating the number of breeding sites. Boxes A, B, and C mark the Iberian Peninsula, Barcelona, and Rome and its surroundings.

**Figure 3.**
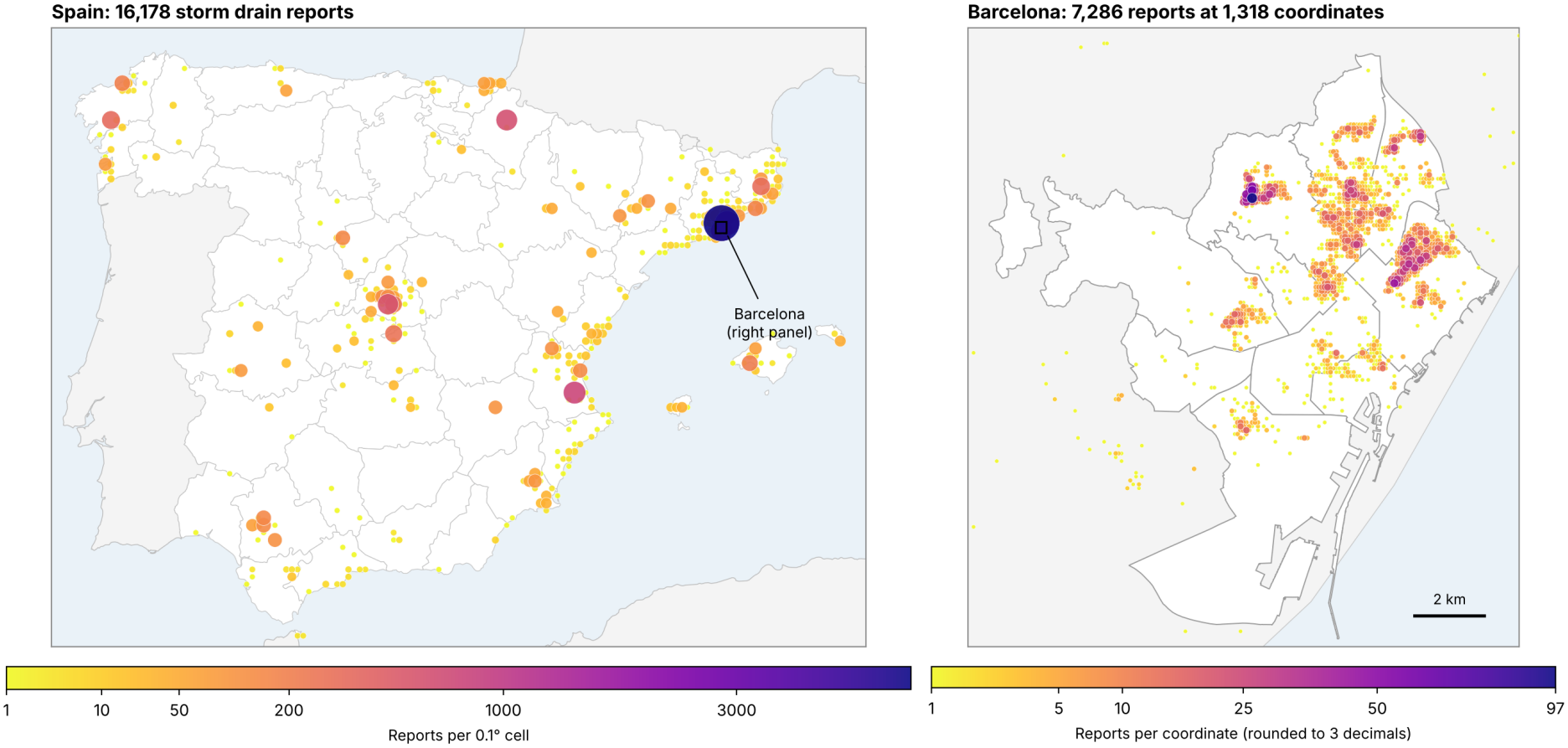
Storm drain reports in Spain (left), counted in 0.1^°^ cells, and a detail of Barcelona (right), counted using coordinates rounded to three decimals, with the boundaries of the city’s ten districts. Circle size and color indicate the number of reports; the two panels use different color scales. One report from the Canary Islands is outside the left panel.

#### Interactive map

We also developed an interactive version of both world maps in Python, exported as a self-contained HTML file. Hovering over a point shows its country, coordinates, and number of breeding sites and, in the water-and-larvae map, how many have water and larvae. Where several coordinates overlap, the tooltip gives the total number of breeding sites, the number of coordinates involved, and the largest one. The map can be zoomed and moved to inspect areas where many coordinates overlap, such as the Iberian Peninsula or Barcelona, and a button switches between the two maps. The interactive map and the script that generates it are available in the repository.

#### Temporal distribution and proportions

Using the local creation time of each report, we counted reports per year and per calendar month (Figure 4), per day, and per day of the week. We then computed the share of storm drains reported with water, for all records and for each country with at least 20 reports, and the share of drains with water in which larvae were observed, overall, for Spain and the rest of the countries, and for the two periods in which larvae were recorded (Figure 5). Each share is shown with a 95% Wilson confidence interval, which describes the sampling uncertainty of a proportion but not the effect of uneven reporting.

**Figure 4.**
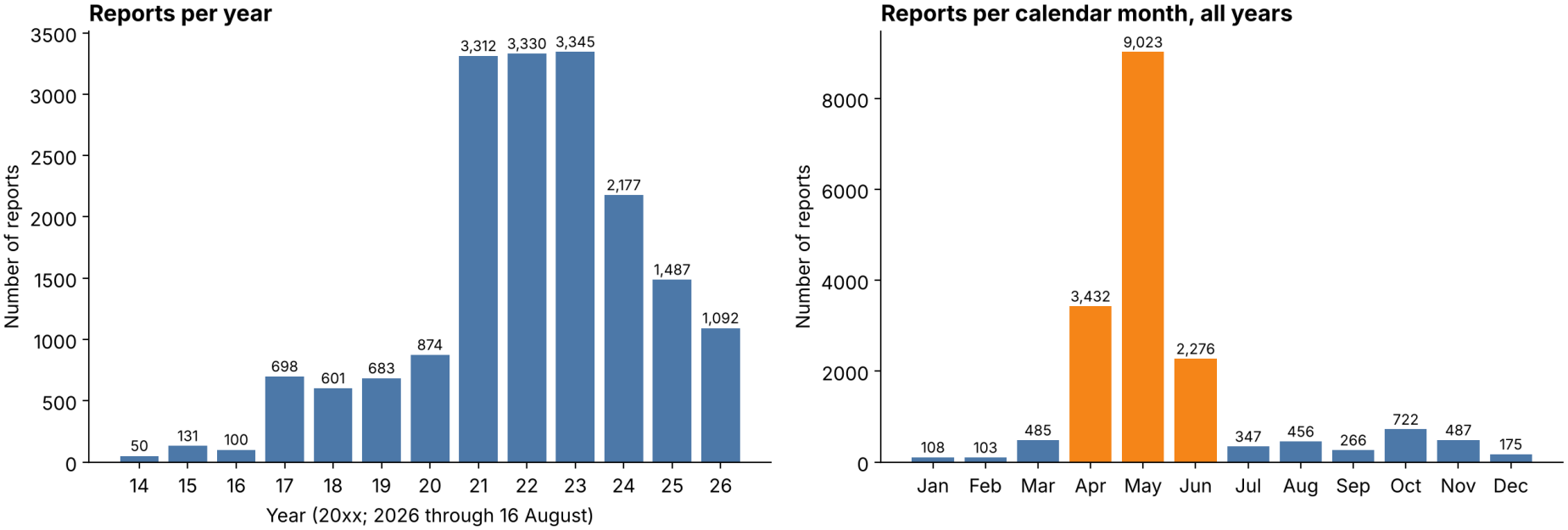
Storm drain reports per year (left) and per calendar month, adding all years (right). The months from April to June are highlighted.

**Figure 5.**
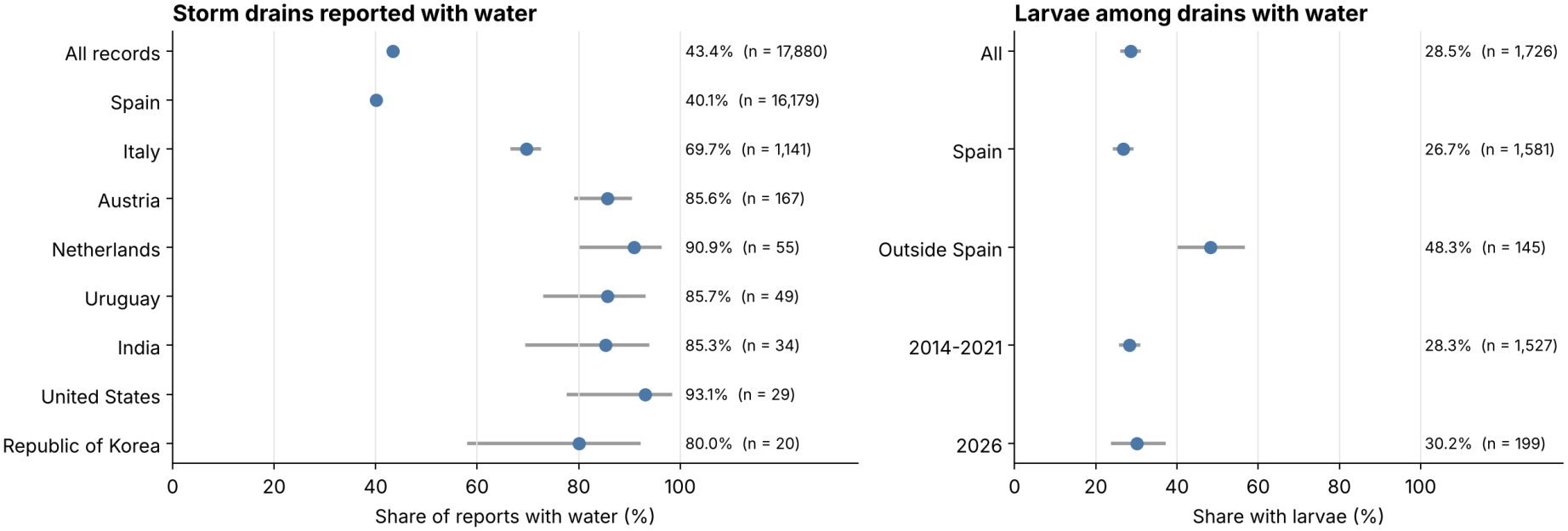
Share of storm drains reported with water, for all records and for countries with at least 20 reports (left), and share of drains with water in which larvae were observed (right). Points are the observed shares and lines are 95% Wilson confidence intervals.

#### Map of Spain and Barcelona

Because most records come from Spain, we also drew a map of that country with a detail of the city of Barcelona (Figure 3). In the map of Spain, reports are counted in 0.1^°^ latitude and longitude cells; in the Barcelona detail, they are counted using coordinates rounded to three decimals, as in the world map. Province and country boundaries come from Natural Earth (1:10m), and the district boundaries of Barcelona from the open data of the Ajuntament de Barcelona. Reports were assigned to districts by testing whether each point falls inside the district polygon.

### 2.3 Group events and participants

To distinguish reports made during collective activities from the rest, we defined a group event as a municipality and local calendar day with reports from at least five different users, and we classified every report as belonging to a group event or not. The dataset does not say how or why a report was made, so this is an operational definition based only on who reported where and when. We repeated the classification with thresholds of three and ten users to check that the results do not depend on choosing five. We then compared the two kinds of reports by day of the week, month, hour, and country, and by the shares of drains with water and with larvae (Figure 6).

**Figure 6.**
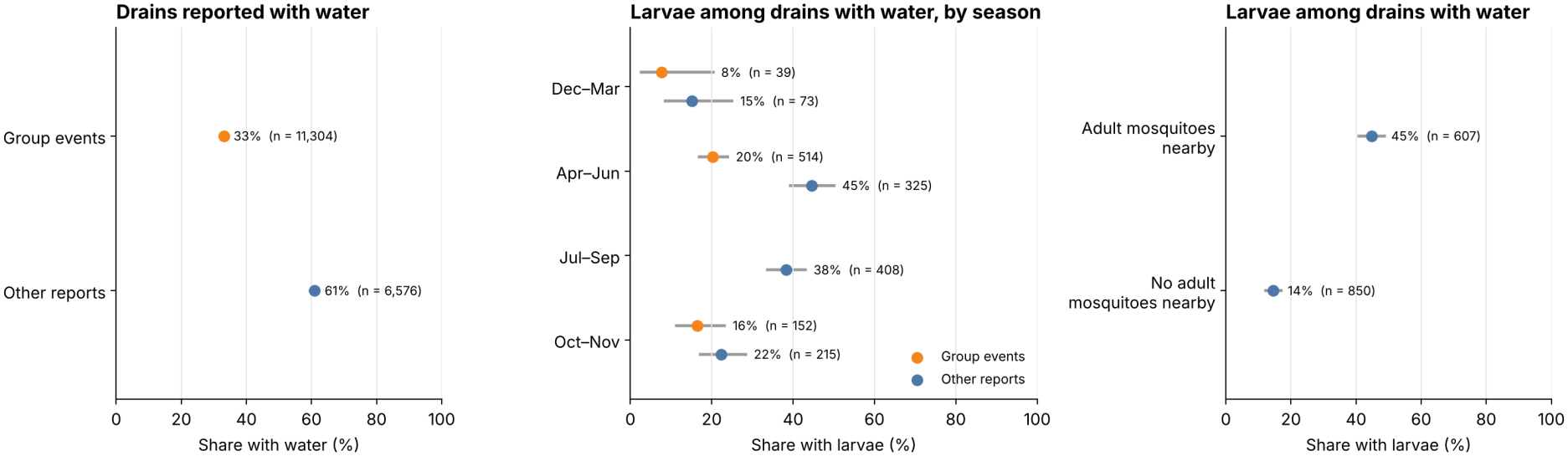
Share of storm drains reported with water in group events and in other reports (left); share of drains with water in which larvae were observed, by period of the year and kind of report (center); and the same share according to whether the participant noticed adult mosquitoes nearby (right). Points are the observed shares and lines are 95% Wilson confidence intervals. No report from a group event between July and September has information on larvae.

**Figure 7.**
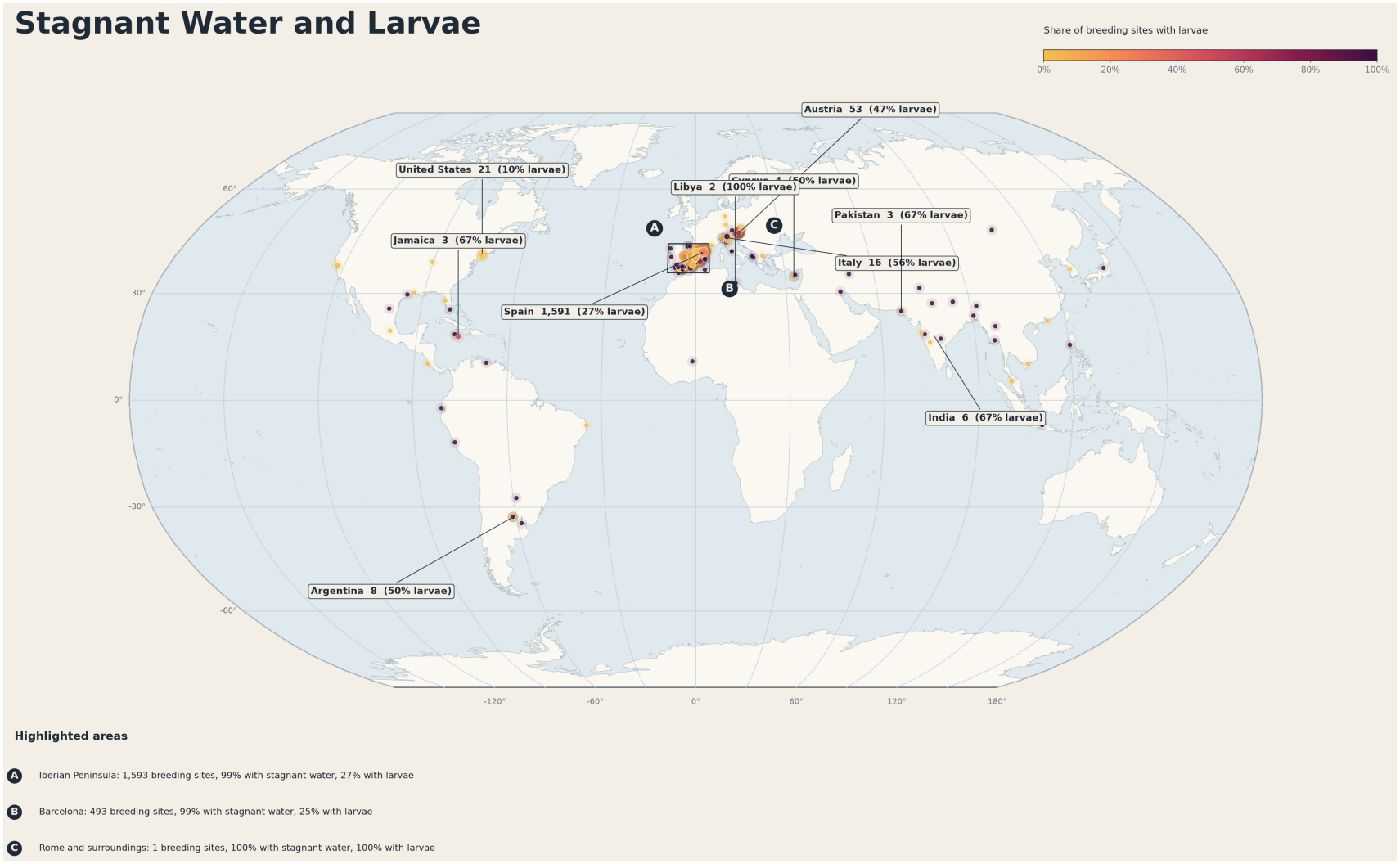
World map of the relationship between stagnant water and larvae in mosquito breeding sites. Circle size indicates the number of breeding sites at each coordinate, and color indicates the share of those sites with larvae

To describe participation, we counted for each user the number of reports, the number of different days with reports, and the number of calendar years with reports. To describe seasonality, we computed the share of drains with water in which larvae were observed in four periods of the year (December–March, April–June, July–September, and October–November), separately for group events and other reports. Finally, we compared the shares of drains with water and with larvae between reports in which the participant noticed adult mosquitoes nearby and reports in which they did not.

### 2.4 Data and Code Availability

The records analyzed in this study were obtained from the Mosquito Alert public map [14], as described in Section 2.1. Because the map is updated continuously, the exact snapshot used in this paper (downloaded on September 12, 2026) is archived together with the code used to produce all figures and the interactive map. The development version is available on GitHub: https://github.com/anaguilar02/mosquito_breedingsites. The interactive map is provided in the repository as the file `breedingsites_map.html`.

## 3 Results

This section presents the results of the mosquito breeding site analysis: frequency by country and geographic coordinates, mapping at global and national scales, distribution over time, group events and participation, and the frequency with which water and larvae are reported.

### 3.1 Where reports are submitted

Figure 1a shows that most breeding sites are located in one country. Spain has 16,179 records, far more than Italy (1,141) and Austria (167), which rank second and third. All other countries have fewer than 60 records each, ranging from 55 in the Netherlands to 5 in the Russian Federation. Within Spain, the records are also concentrated: Catalonia has 9,135, and the municipality of Barcelona alone has 7,217, which is 40% of the whole dataset. The dataset is therefore largely made up of Spanish observations, which should be kept in mind when interpreting the global patterns (Figure 2).

Figure 3 shows this concentration within Spain. Reports come from 44 provinces, but they are grouped around a few places: Barcelona has 8,060 records, followed by Madrid (1,386), Valencia (1,122), Girona (803), Navarre (638), and A Coruña (583). Of the 335 cells with at least one report, four contain half of the Spanish records, and the cell with the most reports, in Barcelona, contains 5,531. After Barcelona, the municipalities with the most reports are Algemesí (769), Madrid (644), and Pamplona (634). The details for Barcelona show that the reports do not cover the city evenly either. They form compact groups of neighboring coordinates, and four of the ten districts, Sant Martí (2,166), Horta-Guinardó (2,152), Nou Barris (1,067), and Sant Andreu (864), contain 87% of the reports located within the city, while large parts of the remaining districts have almost none.

Figure 1b shows the 30 most frequent coordinates in the dataset. Coordinates were rounded to three decimal places, so each bar groups all records within a small area of roughly 100 m, which explains why the counts are much smaller than those by country in Figure 1a. The location (39.909, -3.625) stands out from the rest, with 172 records, nearly twice the next highest value (97 at 41.431, 2.143). After that, the counts gradually decrease from 58 to 28. Of the 30 coordinates, 24 lie in the Barcelona area (latitude 41.41–41.45, longitude 2.14–2.20) and form two neighboring groups, one around longitude 2.14 and another around 2.18–2.20. The remaining six are the top coordinate, in central Spain, two in Mallorca (39.583 and 39.582, 2.638), one in Navarre (42.810, -1.662), one in Murcia (37.976, -1.231) and one in Uruguay (-34.831, -56.218), which is the only site outside Spain.

We inspected the individual reports at the top coordinate, which lies in Yepes (Toledo). Its 172 records were submitted by 52 different users on six days between June 2022 and May 2026, 107 of them on June 9 and 10, 2022. Each report has a different exact position, and 164 of them describe dry drains. The count therefore is not produced by duplicate reports or a default location; it reflects many participants reporting the storm drains of the same small area on the same days. The second coordinate, in Barcelona, shows the same structure: 97 reports from 20 users on seven days.

### 3.2 When reports are submitted

Figure 4 shows that reports are very unevenly distributed in time. Fewer than 900 reports per year were submitted until 2020; the number then rose to about 3,300 per year between 2021 and 2023, which together contain 56% of all records, and has decreased since. Within the year, 82% of all reports were submitted between April and June, and half of them (9,023) in May alone. Reporting is also concentrated on particular days. The dataset covers 1,378 days with at least one report, but the 50 busiest days contain 51% of the records and the 100 busiest contain 68%. On the busiest day, May 27, 2021, 24 users submitted 438 reports. In addition, 90% of reports were submitted from Monday to Friday. Many users on the same weekdays in spring, each reporting many drains in a small area, is the pattern expected from organized group activities rather than from individual, spontaneous reporting.

### 3.3 Group events and participants

The classification into group events supports this reading. There are 165 municipality-days with reports from at least five users, in 53 municipalities, with a median of 8 users and 41 reports each. Together they contain 11,304 records, 63.2% of the dataset. Reports in group events were submitted almost exclusively from Monday to Friday (99.2%), between April and June (92.7%), between 8:00 and 15:00 (91.2%), and in Spain (99.8%). The remaining 6,576 reports are more spread out: 73.3% from Monday to Friday, 64.6% between April and June, 50.9% between 8:00 and 15:00, and 74.5% in Spain. Group events contributed 15% or less of the reports of each year until 2016 and between 32% and 90% from 2017 on. The definition is conservative, because it leaves out smaller groups and single users who report many drains in one day. With thresholds of three and ten users, group events contain 71% and 45% of the records, respectively, and the differences in water described below remain (35% against 65% with three users, and 30% against 54% with ten).

Participation is also brief. Of the 3,313 users, 87.7% submitted all their reports on a single day and only 1.4% reported in more than one calendar year; 41.6% submitted a single report, and the median is two. About half of the users (48.0%) took part in at least one group event. Reports are concentrated in a few participants: the most active 10% of users submitted 47.7% of the reports.

### 3.4 Water and larvae

Of the 17,880 storm drains, 7,756 (43.4%) were reported with water and 10,124 as dry. In absolute numbers, the drains with water follow the same ranking as the reports: 6,485 are in Spain, 795 in Italy, and 143 in Austria, and every other country has 50 or fewer.

The share of drains reported with water gives a different picture (Figure 5, left): Spain has the lowest share, 40.1% (95% CI 39.3–40.8), compared with 69.7% (66.9–72.3) in Italy and 80% or more in the other countries with at least 20 reports. In other words, most dry storm drains in the dataset were reported in Spain. Within Spain, the share with water was 35.2% among the 13,747 reports submitted between April and June and 67.9% among the 2,432 reports submitted in the rest of the year.

Group events explain much of these differences (Figure 6, left). Water was reported in 33.2% (95% CI 32.3–34.1) of the drains in group events and in 60.9% (59.7–62.0) of the drains in other reports. Leaving out group events, the share with water in Spain rises to 56.2% (2,756 of 4,902), closer to the 74.7% (1,271 of 1,701) observed in the rest of the countries.

Figure 1c shows the number of breeding sites with and without larvae, depending on whether stagnant water was present. Almost all the records in this figure have stagnant water: 1,726 of the 1,739 sites (1,234 without larvae and 492 with larvae), while only 13 sites have no stagnant water (eight without larvae and five with larvae). This does not mean that almost all storm drains hold water; it reflects that larvae were recorded almost only for drains with water (Table 1). Among the 1,726 drains with water and information on larvae, larvae were observed in 492 (28.5% [95% CI 26.4–30.7]), so sites without larvae are about 2.5 times more frequent than sites with larvae (Figure 5, right). The share was similar in the two periods with data, 28.3% until 2021 and 30.2% in 2026. It was lower in Spain, 26.7% (24.6–28.9; 422 of 1,581), than in the rest of the countries taken together, 48.3% (40.3–56.3; 70 of 145). The sites without stagnant water are too few to compare with the rest.

The share of drains with larvae also depends on the kind of report and on the time of year (Figure 6, center). Larvae were observed in 18.7% (16.0–21.8; 132 of 705) of the drains with water reported in group events and in 35.3% (32.4–38.2; 360 of 1,021) of those in other reports. By period of the year, the share was 12.5% from December to March, 29.7% from April to June, 38.2% from July to September, and 19.9% in October and November. Because group events are concentrated in spring, the seasonal comparison is clearer within the other reports: 15.1% (11 of 73) from December to March, 44.6% (145 of 325) from April to June, 38.2% (156 of 408) from July to September, and 22.3% (48 of 215) in October and November. In these reports, larvae were therefore observed in about 40% of the drains with water between April and September and in about 20% during the rest of the year.

Finally, the reports are internally consistent (Figure 6, right). Among the 2,502 reports that record whether adult mosquitoes were noticed nearby, water was reported in 76.6% of the drains with mosquitoes nearby and in 49.7% of the drains without them. Among the drains with water and information on larvae, larvae were observed in 44.8% (40.9–48.8; 272 of 607) when adult mosquitoes were noticed nearby and in 14.5% (12.3–17.0; 123 of 850) when they were not. The difference appears both in group events (31.1% against 13.9%) and in other reports (50.6% against 15.4%).

Figure 7 shows the geographic distribution of the 1,739 breeding sites with information on both stagnant water and larvae. Overall, larvae were found in 497 of these sites. The map is dominated by Spain, which has 1,591 of these sites and a 27% share of larvae, very close to the overall value. Within the highlighted areas, the Iberian Peninsula (A) has 1,593 sites, 99% with stagnant water and 27% with larvae, and Barcelona (B), within area A, has 493 sites, with 99% and 25%, respectively. Rome and its surroundings (C) have only one record, so its percentages do not represent a pattern. Outside Spain, Austria, the United States, and Italy have the most records, and the remaining labeled countries have between two and eight sites each. Since almost all sites in this subset have stagnant water, the map color mainly reflects the presence of larvae. However, most coordinates outside Spain have only one or a few records, so their color is usually at one of the extremes and should not be interpreted as a trend.

## 4 Discussion and Conclusions

In this work, we explored the storm drain breeding sites reported through Mosquito Alert from June 2014 to August 2026, describing their distribution by country, coordinate, and time, the frequency of stagnant water, and its relationship with the presence of larvae. Since the analysis is descriptive, the following interpretations are preliminary and should be treated as hypotheses for future work.

The most evident feature of the data is its concentration in space and in time. Spain accounts for 16,179 of the 17,880 records (about 90%), the municipality of Barcelona for 40% (mostly in four of its ten districts), and 24 of the 30 most frequent coordinates are in the Barcelona area. In time, 82% of the reports were submitted between April and June, and half of all records fall on 50 days. This suggests that the number of reports reflects where and when the application is used rather than where storm drains or mosquitoes are most abundant, in line with the spatial sampling biases described for citizen science data [15]. The reports at the most frequent coordinate (39.909, -3.625; 172 records) illustrate this: they are not repeated reports but the work of 52 users on a few days. More generally, 63% of the records were submitted on days when at least five users reported in the same municipality, almost always on weekdays, during the morning and early afternoon, and in spring. This pattern is expected from organized activities, such as educational or community campaigns, although the dataset does not identify them and their nature would need to be confirmed with the platform. Most participants, in turn, reported on a single day. For these reasons, the comparisons between countries or cities in this study should not be interpreted as differences in mosquito presence, and the seasonal profile of the reports should not be interpreted as the seasonality of mosquito activity.

The same structure applies to the water information. Overall, 43% of the reported storm drains contained water, but the share was 33% in group events and 61% in the remaining reports, and this difference accounts for much of the gap between Spain (40%) and the other countries (70% or more). A plausible explanation is that participants in organized activities report every drain they find, dry or not, whereas individual users mostly report drains that catch their attention because they hold water; differences in rainfall between seasons and countries may also contribute. The share of drains with water therefore measures how the reports were collected as much as the state of the drains.

Larvae could be studied only in 1,726 drains with water, reported until 2020 or from late May 2026, because the field is missing for the 13,651 reports from 2021 to 2025 and for almost all dry drains. In this subset, larvae were reported in 28.5% of the drains, so most storm drains with water had no visible larvae at the time of the report. This does not mean that larvae were absent, since a report is a single observation by a volunteer at one moment. This share also depends on how the report was collected: it was 19% in group events and 35% in the remaining reports, so the overall value is a mixture of the two. The share was similar in Spain (26.7%) and in the Barcelona area (25%), and higher in the reports from outside Spain taken together (48.3%). With only 145 such reports from many countries, and given the differences in how reports are collected, this difference should not be read as a difference in larval presence between regions. Two results nevertheless suggest that the reports contain information about mosquitoes and not only about reporting. First, outside group events, larvae were observed about twice as often between April and September as during the rest of the year, which agrees with the warm-season activity of *Aedes* mosquitoes in temperate regions. Second, larvae were reported three times as often when the participant also noticed adult mosquitoes nearby, in both kinds of reports. Both results rest on a few hundred participant observations, without expert validation, and adult mosquitoes near a drain need not come from it, but they suggest that individual reports of storm drains with water are a reasonable starting point for locating productive breeding sites.

These results suggest four lessons for using citizen science reports in larval surveillance, relevant wherever platforms of this kind are considered a complement to routine entomological surveillance, including dengue-endemic settings such as Costa Rica. First, report counts measure reporting effort, so they should be interpreted as coverage; statements about the drains require proportions with explicit denominators, and even these depend on how the reports were collected. Second, the way a report was collected should be recorded with it, or at least reconstructed as we did here, and reports from collective activities and from individual users should be analyzed separately, because pooled shares of water and larvae are mixtures of two different sampling processes. Third, the two kinds of reports are complementary rather than interchangeable. Collective activities document many drains in a small area, including dry ones, and provide the denominators that individual reporting lacks; if repeated in the same areas at different times of the year, they could support estimates of how often drains hold water and larvae. Individual reports, in contrast, are selective but more often point to drains with water and larvae. Fourth, the fields that carry the entomological information need to be recorded consistently. Here, the presence of larvae is missing for the five years with the most reports, and a missing value cannot be distinguished from larvae that were not looked for. Since most participants report on a single day, sustained coverage is also more likely to come from recurring activities than from the retention of individual users.

Overall, the dataset is useful for describing where and when reports are submitted, but it is not enough to draw conclusions about mosquito distribution. The main limitations are the uneven reporting effort in space and time, the absence of information on larvae for 90% of the records, including the years with the most reports, the fact that water and larvae are reported by participants without expert validation, the operational definition of group events, which depends on a threshold and cannot be checked against a record of actual campaigns, and the rounding of coordinates to three decimals, which groups records within roughly 100m. Future work could normalize the reports by population or number of active participants, identify the organized activities behind the group events, compare the reports with traditional surveillance data, and analyze their temporal patterns once reporting effort is accounted for, for example with wavelet methods such as those used to study seasonality in dengue incidence [3]. The interactive map can support this process by making it easier to inspect areas where many records overlap.

## Ethics Statement

This study used only openly available, anonymized records from the Mosquito Alert public map [14]. The records identify participants only by an anonymous code, and the analysis reports aggregate results. The study involved no intervention or interaction with participants and no access to identifiable private information, so the authors did not seek ethics committee approval.

## Use of Artificial Intelligence

During the preparation of this work, the authors used Claude (Anthropic), a generative artificial intelligence assistant. It was used to format the manuscript, to review it for errors and inconsistencies, and to revise its wording; to write and run the Python code for the analyses of field completeness, temporal distribution, proportions with confidence intervals, group events, and participants, and for the map of Spain and Barcelona, including the corresponding figures; to draft the passages of the Methodology, Results, and Discussion that describe those analyses; and to search the literature on the validation of Mosquito Alert reports. The study design, data collection, remaining figures, and interactive map are the work of the authors. The authors reviewed and edited all content produced with the tool, verified the results, and take full responsibility for the content of this article.

## Acknowledgments

We thank the Mosquito Alert project and its participants for making their data openly available. F.S. and A.A.P. extend their gratitude to the Research Center in Pure and Applied Mathematics and the Department of Mathematics at the University of Costa Rica for their support.

